# Use of the non-paretic arm reflects a habitual behaviour in chronic stroke

**DOI:** 10.1101/2024.09.09.611968

**Authors:** S. Sporn, E. Bonyadin, R. Fathana, L. Triccas Tedesco, M. Coll, S. Bestmann, N. S. Ward

## Abstract

**Background:** A proportion of stroke survivors use their affected arm less than might be expected based on their level of impairment. The resulting non-use of the affected arm has a negative impact on participation in neurorehabilitation and functional independence. However, non-use remains poorly understood. One possibility is that prioritising the non-paretic arm reflects a habit, despite residual functional capacity in the paretic arm.

**Methods:** 30 chronic stroke survivors (Mean FM: 28.9 ± 11.3) participated in a simplified version of the forced response paradigm, which reliably identifies the presence of a habit. Participants were asked to choose which arm to use to maximise points scored during a reaching task. During half of the trials, the presumed habit of using the non-paretic arm yielded more points, whereas in the other half using the non-paretic arm incurred a loss of points. Participants completed two versions of this task, once with unlimited response time available and once without.

**Results:** Participants scored fewer points in the limited response condition compared to the unlimited response conditions. This difference was driven by a selective increase in the use of the non-paretic arm in trials where the paretic arm yielded more points. The results were not mediated by former hand dominance.

**Conclusions:** Our results demonstrate that not using the non-paretic arm may reflect a habit response that is more readily triggered in demanding (e.g. time-limited) situations. This may explain why successful neurorehabilitation does not always result in a more functionally useful arm. Our results pave the way for targeted interventions such as habit breaking techniques to be included in clinical practise.

## Introduction

Neurological conditions are the leading cause of overall disease burden globally with stroke being the major contributor (Steinmetz et al., 2024). Amongst people with stroke, upper limb impairment is a common contributor to disability, occurring in approximately 75% of cases (Broeks et al., 1999; Lawrence et al., 2001; Nakayama et al., 1994). Some hemiparetic stroke survivors incorporate their paretic affected arm in daily activities less than would be expected based on their impairment (Andrews & Steward, 1979; Sterr et al., 2002), with detrimental effects on their recovery (Buxbaum et al., 2020; Ballester et al., 2022). Identifying factors that contribute to lower than expected levels of paretic arm use will help identify much needed therapeutic targets.

One possibility is that prioritising the non-paretic arm during daily activity reflects habitual behaviour. Habits are automatic responses that can be differentiated from goal-directed responses (Figure 1a). Habitual responses are fast, yet inflexible, while goal-directed responses are slow, yet highly flexible (Robbins and Costa, 2017; Doan and Dayan, 2013; Wood and Rünger, 2016). Habitual responses are useful, because they are fast at triggering actions that have led to the most rewarding outcomes in the past. Yet, when reward contingencies change, habitual responses can lead to inefficient behaviour. Immediately after hemiparetic stroke for example, patients may prioritise use of the non-paretic arm because it helps them achieve immediate goals (e.g. dressing, feeding etc.). However, as some recovery occurs and more paretic arm use is possible, patients may continue to prioritise non-paretic arm use (for immediate goal attainment), to the detriment of the longer-term aim of functional recovery of the paretic arm. Here, we tested the hypothesis that prioritising the non-paretic arm during activities may reflect a habitual response in chronic stroke patients.

**Figure 1.**
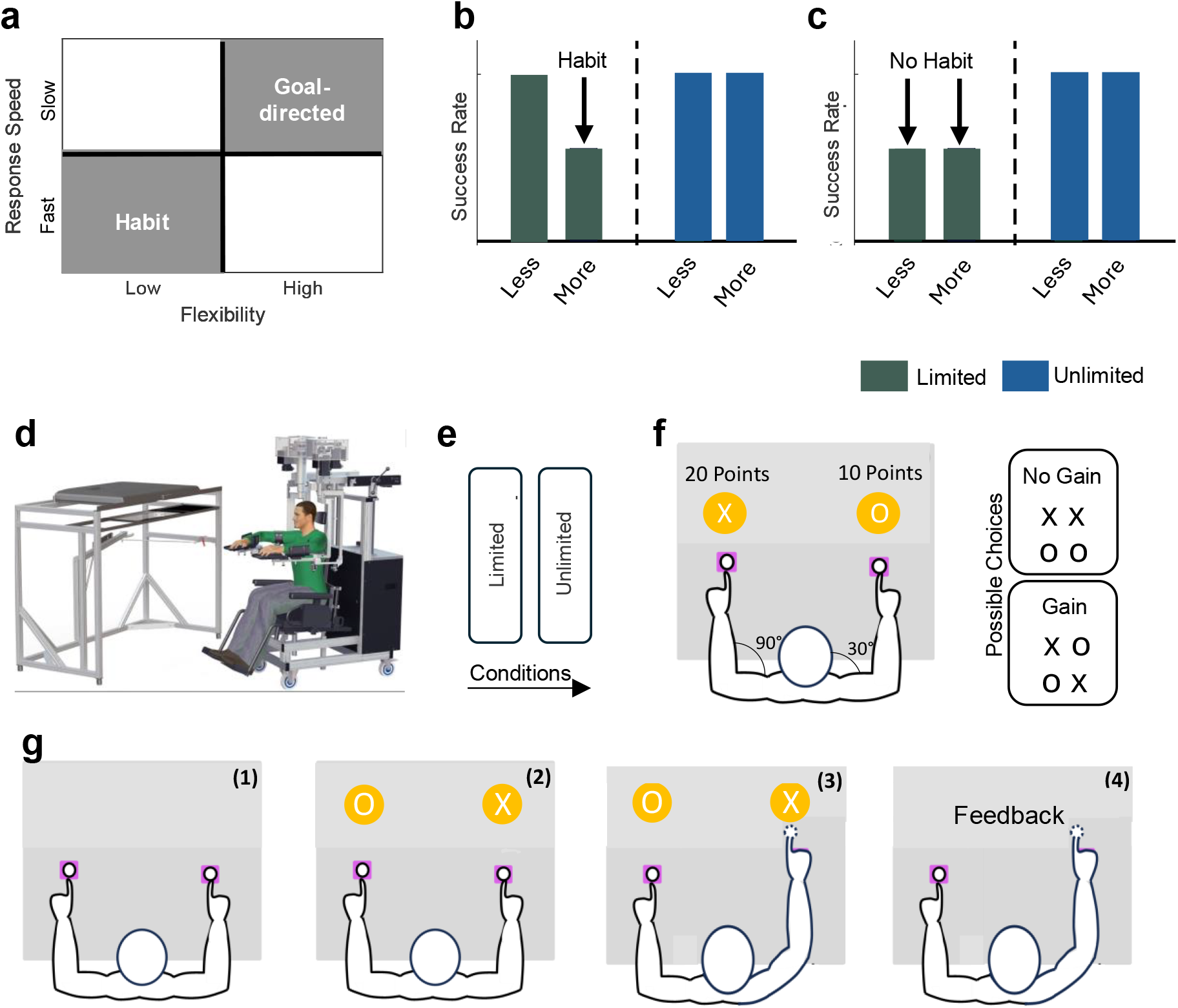
Theoretical Background and Experimental Design. **a)** Habitual and goal-directed responses have orthogonal properties. While habitual responses are fast, yet inflexible, goal-directed response s are slow, yet flexible. **b)** A habit is present if *Success Rate* decreases in trials with limited response times available where the habitual response does not lead to the most rewarding outcome (*More/Limited*). **C)** In contrast, a decrease in both the *Less* and *More* condition suggests the opposite. **d)** Illustration of the KINARM Exoskeleton; a robot which gathers arm kinematic data during task performance. **e)** Experimental phases. *Unlimited response* phase: Participants have unlimited deliberation time; *Limited response* phase: Participants are instructed that they must make fast responses. **f)** Illustration of the workspace and of the arm configuration. Participants were asked to make 2cm reaching movements from a starting position to a peripheral target. Both arms were aligned so that the starting positions were at the midpoint of each arm’s workspace (90° elbow flexion and 30° shoulder flexion. **g)** Illustration of a single trial. **1)** Participants were asked to move their cursors into the starting positions and wait for one of the possible choices to appear (300-350ms). **2)** Participants were told each target would contain either the symbol X, denoting 20 points, or O, denoting 10 points. **3)** Participants had to select one target by reaching into it, and were instructed to try to earn as many points as possible. **4)** After reaching into a target, text feedback stating how many points had been earned in this trial appeared above the targets for 800ms. Participants then returned to the starting positions for the next trial. Participants completed 48 trials in both phases.

Habitual behaviour after stroke can be identified with the *forced response paradigm (*Keramati et al., 2011; Hardwick et al., 2019; Luque et al., 2020). Participants are asked to choose which arm to use during a reaching task in which the goal is to maximise reward. For half the trials using the paretic arm is associated with the highest reward (paretic trials), while for the other half using the non-paretic arm is associated with the highest reward (non-paretic trials). Choosing the arm that is associated with less reward in either trial is considered an error. In addition, the task is performed under two conditions, with either unlimited or limited time to make a response (‘forced’).

In the unlimited response time condition, participants should be able to maximise points with few errors by choosing the arm that yields more points (Figure 1a, b). It is during the unlimited response time condition that the presence of a habitual response can be detected. During non-paretic trials, subjects only have to use the non-paretic arm to maximise reward. However, during paretic, high reward will come with paretic arm use, but a habitual response with the non-paretic arm (brought on by limited available response time) will result in reduced success rate for in this trial condition only (Figure 1a). If however, there is no habitual response, we would expect the time-limited condition to lead to a similar increase in errors for paretic and non-paretic arm use (i.e., random selection of either arm; Figure 1b).

## Methods

### Patient recruitment

30 chronic stroke patients (≥ 6 months from stroke onset) were recruited from the Queen Square Upper Limb rehabilitation programme (QSUL) for the experiment. The inclusion criteria for this experiment were: 1) first ever stroke and 2) no other brain injury, neurological condition or major psychiatric illness, while exclusion criteria were: 1) hemi-spatial neglect or hemianopia, 2) severe aphasia, or 3) pain limiting ability to participate in tasks or follow the study protocol. All participants were comprehensively informed about the study, and written consent was obtained before their participation.

### Experimental apparatus

The study utilized the KINARM Exoskeleton (BKIN Technologies Ltd, Kingston, ON, Canada; Figure 1d). This robotic apparatus collects kinematic data of the arms during various tasks. Designed to support the arms, forearms, and hands of the user, the KINARM Exoskeleton permits only horizontal movements, primarily involving the flexion and extension of the shoulder and elbow joints. Participants are seated with their arms outstretched horizontally, typically at an 85-90 degree angle from their shoulders. Each arm segment is equipped with a customized exoskeleton for enhanced comfort and support. Additionally, the device is integrated with a 2-D virtual reality display, aligning with the arms’ plane, to provide visual cues and feedback. Calibration is conducted prior to each session to ensure precise tracking and interaction within the virtual setup. Notably, while the robot provides gravitational support, it does not aid in the completion of tasks within the experiment. Further details about the KINARM Exoskeleton can be found in the works of Scott (1999), Coderre et al. (2010), and Dukelow et al., (2010).

### Experimental design

To test the hypothesis that non-use reflects a habitual response in chronic stroke we modified an established paradigm to elicit habitual responses (Keramati et al., 2011; Hardwick et al., 2019; Luque et al., 2020). It represents a simplified version of the *forced response paradigm* (Figure 1e; Keramati et al., 2011; Hardwick et al., 2019; Luque et al., 2020).

In the present experiment, participants chose between two letters (‘X’ and ‘O’) that were associated with 20 points (‘X’) and 10 points (‘O’), respectively. In each trial, one of the four possible combinations (i.e., X–X, O–O, X–O, O–X, Figure 1f) was presented. Participants had to decide which arm to use, to maximise reward. Importantly, the left target could only be reached for with the left arm, while the right arm could only aim for the right target (Figure 1f). Therefore, during *Gain* trials (i.e., X–O, O–X), participants could maximise reward by selecting the arm that yields more points. At times the more rewarding option corresponded to choosing the non-paretic arm. Therefore, here the hypothesised habit of prioritising the less arm aligns with the choice that maximises points (non-paretic trials). More importantly, however, in the other *Gain* combination using the non-paretic arm represented the less rewarding option. Therefore, here the using the non-paretic arm does not align with the goal of the task (paretic trials; Figure 1f). In *Neutral* trials (X-X and O-O) choosing either arm returned the same number of points. All four combinations were randomly presented in one block (12 blocks in total).

During each trial, two purple starting positions with a radius of 2cm appeared which participants were instructed to return to between trials. Each starting position was positioned so that the shoulder and the elbow were at approximately 30° and 90°, respectively (Figure 1f). The starting positions were, therefore, level on the y-axis, with one on each side. Participants were instructed to reach into each circle only with the ipsilateral hand. After maintaining position in the starting position between 300-350ms (with timings randomly varied to prevent anticipatory movements; Figure 1g, 1)), two orange targets (radius of 3cm) appeared 2cm directly above the starting positions containing either an X or O (one above each starting position; Figure 1g, 2)). The symbols X and O were chosen to reduce demands on mathematical ability. Participants had to select one target by reaching into it, and were instructed to try to earn as many points as possible. Participants did not have to stay in the targets, with overshoot allowed to reduce overall accuracy demands. The cursors were visible throughout this task (Figure 1g, 3)). Immediately after reaching into a target, feedback (in white text) stating how many points had been earned in this trial appeared above the targets for 800ms (Figure 1g, 4)). Participants then returned to the starting positions for the next trial Participants were asked to complete the experiment twice (Figure 1e). During the *Unlimited* condition participants had unlimited time to respond. However, during the *Limited* condition response times were restricted and participants were instructed that decisions about arm choice had to be made fast. Specifically, participants were informed that long response times triggered failed trials which yielded 0 points.

Participants completed the task twice: once with unlimited response time available and once without. In summary, participants engaged in 4 trial conditions: 1) *Non-Paretic/Unlimited* where choosing the non-paretic arm maximised points while having unlimited time to respond; 2) *Paretic /Unlimited* where choosing the paretic arm maximised points while having unlimited time to respond 3) *Non-Paretic/Limited* where choosing the non-paretic arm maximised points while having to respond; and 4) *Paretic /Limited* where choosing the paretic arm maximised points while having to respond.

During the *Limited* condition participants completed 48 trials, corresponding to 6 blocks of 8 trials with all 4 trade-off combinations included twice. Importantly, unbeknownst to the participants, in each block one set of all possible combinations triggered a failed trial irrespective of how fast they responded in these trials. These failed trials were included to reinforce that participants were required to respond fast in order to trigger habitual responses (Hardwick et al., 2019). This simplified version of the *forced response paradigm* was favoured over the original design (Hardwick et al., 2019) because of several reasons. In the original design participants were presented with a sequence of four auditory stimuli, with an inter-stimulus interval of 400 milliseconds, and were tasked to time their response to coincide with the fourth and final tone. The timing of the stimulus presentation was randomised, chosen from a uniform distribution across the sequence, to systematically modulate the preparatory time available to the participant to respond. Responses that were triggered 100ms before or after the final tone were deemed failed trials. Despite its elegance, we reasoned that this design requires a substantial amount of cognitive resources (i.e., sustained attention) which may challenge even mildly cognitively impaired stroke survivors (Spaccavento et al., 2019; Barker-Colo et al., 2010; Hyndman and Ashburn, 2003). Additionally, 500 *limited response time* trials were conducted in the original design to cover the whole preparatory time window (0-1200ms) which was deemed unfeasible for a study involving chronic stroke survivors. Finally, a fixed cut-off of 100ms may render a high number of trials as failed trials especially considering that processing speed and cognitive fluidity may be very heterogeneous in a group of chronic stroke survivors. The aim of the *Limited* condition is to enforce faster decision-making which may lead to habitual responses being triggered. We, therefore, included a number of failed trials to reinforce that participants made fast decisions, which was the same across participants. Participants received a feedback note in red stating that the participants should ‘Try to make faster decisions – 0 points’ after they chose which arm to use.

#### Trail Making Test

In between the *Unlimited* and *Limited* condition participants were asked to complete the standardised KINARM task *Trail Making Task* (TMT) with their non-paretic arm. The TMT (Part A) was administered to test participants’ cognitive abilities (Bowie & Harvey, 2006), to investigate if performance on this task is a function of cognitive impairment. This is a test of visual search and motor speed skills, and is believed to be sensitive to cognitive impairment in those with neurological conditions (Bowie & Harvey, 2006). Importantly, it is also believed to be a reliable measure of processing speed (Bowie & Harvey, 2006), which is important in making goal-directed decisions (Schad et al., 2014), and fluid cognitive ability (Salthouse, 2011). Participants were instructed to connect 25 semi-randomly dispersed numbered targets in ascending order as ‘fast and accurately as possible’. All targets were presented simultaneously. A cursor followed participants’ hands. If participants hit an incorrect target, the target they had successfully reached before turned red (from white) and participants had to return to that target before proceeding. Before each 1-25 run, participants completed a practice run with five targets numbered 1-5. No time limits were imposed.

### Outcome variables

The 2D (x, y) position of the index finger was recorded at 1000Hz by the KINARM Exoskeleton and was analysed ‘offline’ using Matlab (version R2019b, The MathWorks, Natick, MA, USA).

#### Success Rate

*Success Rate* (in %) reflects how often participants chose the arm that maximised points in the *Unlimited* and *Limited* condition, respectively. Only *Gain* trials were included in this analysis (i.e., *Non-Paretic* and *Paretic* trials).

#### Non-Paretic Arm Use

*Non-Paretic Arm Use* reflects how often the non-paretic arm was chosen (in %), while its inverse indicates how often the paretic arm was chosen (*Paretic Arm Use*). This was based on which arm successfully hit the target. However, trial-by-trial analysis of the velocity profiles of both arms indicated that in 38 trials (1.43%) a reaching movement with the non-successful arm (i.e., the one that did not reach the target) preceded the successful reaching movement. *Non-paretic Arm Choice* was corrected in these trials to accurately reflect arm choice. Additionally, trials in 9 (0.34%) trials both arms were used which were subsequently excluded from further analysis.

#### Dominant Arm Use

*Dominant Arm Use* reflects how often the dominant arm was chosen (in %), while its inverse indicates how often non-dominant arm was chosen.

#### Response Times

Conceptually, *Response Times* (in seconds) were thought to represent the time of movement onset. Movement onset was determined using a weighted measures approach that included 3 trial-based vectors: 1) velocity, 2) acceleration and 3) time passed. In this analysis movement onset is characterised by low values in both the velocity and acceleration vector and higher values in time passed (i.e., movement onset should occur after the presentation of the choice). The aim of this analysis was to find the minimum value of the sum of all three vectors, which is thought to best fit with the time of movement onset. Based on previous research demonstrating that RTs < 300ms reflect random choices (neither goal-directed not habitual), trials with RTs < 300ms were excluded from further analysis (Hardwick at al., 2019). This amounted to 89 trials (3.35%). Medians were used because one-sample Kolmogorov-Smirnov tests (`kstest`) indicated that the data was not normally distributed.

### Analysis plan

#### 1) Do chronic stroke patients exhibit a habit of using the non-paretic arm?

A decrease in *Success Rate* only in the *Paretic/Limited* trial condition suggests a shift from triggering goal-directed to habitual responses, because here the habitual response corresponds to using the non-paretic arm more despite this not being the more rewarding choice. To this end, we assessed changes in use of the non-paretic arm in 3 ‘Choice Conditions’: 1) *Paretic*. Here, using the non-paretic arm would result in a loss of 10 points, while in 2), *Non-Paretic* selecting the non-paretic arm yields more reward. In contrast, in 3) *Neutral* using either arm results in the same number of points (e.g., X-X and O-O). A repeated-measures ANOVA was conducted in MATLAB with ‘Response Condition’ (*Unlimited* vs *Limited* response time) and ‘Choice Condition’ (*Paretic, Non-Paretic* and *Neutral*) as within factors and *Non-Paretic Arm Use* as the dependent variable. Post-hoc analysis included independent Wilcoxon Rank Sum Tests which were corrected for multiple comparisons using Bonferroni Corrections while Cohen’s d was used to estimate effects sizes. A significant increase in *Non-paretic Arm Use* in the *Paretic /Limited* condition would indicate that participants increase the use of their non-paretic arm despite this being the less rewarding choice (habitual response).

#### 2) Is the habit response related to changes in the use of dominant arm?

To investigate if *Dominant Arm Use* affects arm use from the *Unlimited* to the *Limited* response time condition, we used the same repeated-measures ANOVA analysis pipeline as above with DC as the dependent variable. A lack of change in dominant arm use in the *Paretic/Limited* condition would indicate that former arm dominance does not affect arm use across phases.

#### 3) Did participants Response Times decrease from the Unlimited to the Limited response phase?

To investigate if *Response Times* decrease from the *Unlimited* to the *Limited* response time condition, we used the same repeated-measures ANOVA analysis pipeline as above with *Response Times* as the dependent variable. A significant decrease in *Response Times* across phases in all conditions would indicate that deliberation times are significantly reduced and suggests that this simplified version of the *forced response* paradigm worked.

#### 4) Are changes in Non-paretic Arm Choice related to motor and/or cognitive impairment?

To assess if motor and/or cognitive impairment modulates arm choice, we ran independent correlations between changes in the use of the paretic arm (*Paretic Arm Use*) across Response Conditions for both the *Paretic* and *Neutral* conditions and FMA scores (FMA Shoulder and FMA Elbow) and TMT scores. A lack of a significant correlation for any contrast would indicate that changes in *Paretic Arm Use* are independent from motor and/or cognitive impairment and suggests that non-use is an independent therapeutic goal.

## Results

### Study population

30 chronic stroke patients (≥ 6 months from stroke onset) admitted to the Queen Square Upper Limb rehabilitation programme (QSUL) were recruited for this experiment. The clinical and demographic characteristics are summarised in Table 1.

**Table 1.** Clinical and demographic characteristics participants.

Characteristic
|  |  |
| --- | --- |
| Gender – male: female | 20:10 |
| Handedness (prior to stroke) – right: left | 23:7 |
| Affected upper limb – dominant: non-dominant | 20:10 |
| Age (years), mean (SD) | 51.40 (16.8) |
| Time since stroke (months), mean (SD) | 34.7 (37.2) |
| Fugl Meyer Assessment Upper Limb (FMA-UL) (total 54 points), mean (SD) | 28.9 (11.3) |

#### 1. Increased use of non-paretic arm despite it being the less rewarding choice

Success rates were only reduced in the *Paretic/Limited* condition (Figure 2a) suggesting that participants expressed a habit (as seen in Figure 1b). Results from the repeated-measures ANOVA revealed a significant main effect for ‘Response Condition’ (F = 36.54, p < 0.0001, η^2^ = 0.20, Figure 2b) and ‘Choice Condition’ (F = 802.61, p < 0.0001, η^2^ = 0.97). Importantly, we also found a significant interaction between ‘Response x Choice Condition’ (F = 30.52, p < 0.0001, η^2^ = 0.29). Post-hoc pairwise comparisons revealed that *Non-paretic Arm Use* significantly increased in the *Limited* Response compared to the *Unlimited* Response Condition in *Paretic*(Z = 3.55, p = 0.0008, d = 0.97, Figure 2c) and *Neutral* (Z = 3.91, p = 0.0001, d = 0.66), but not in *Non-Paretic* (Z = -1.64, p = 0.1007, d = -0.41).

**Figure 2.**
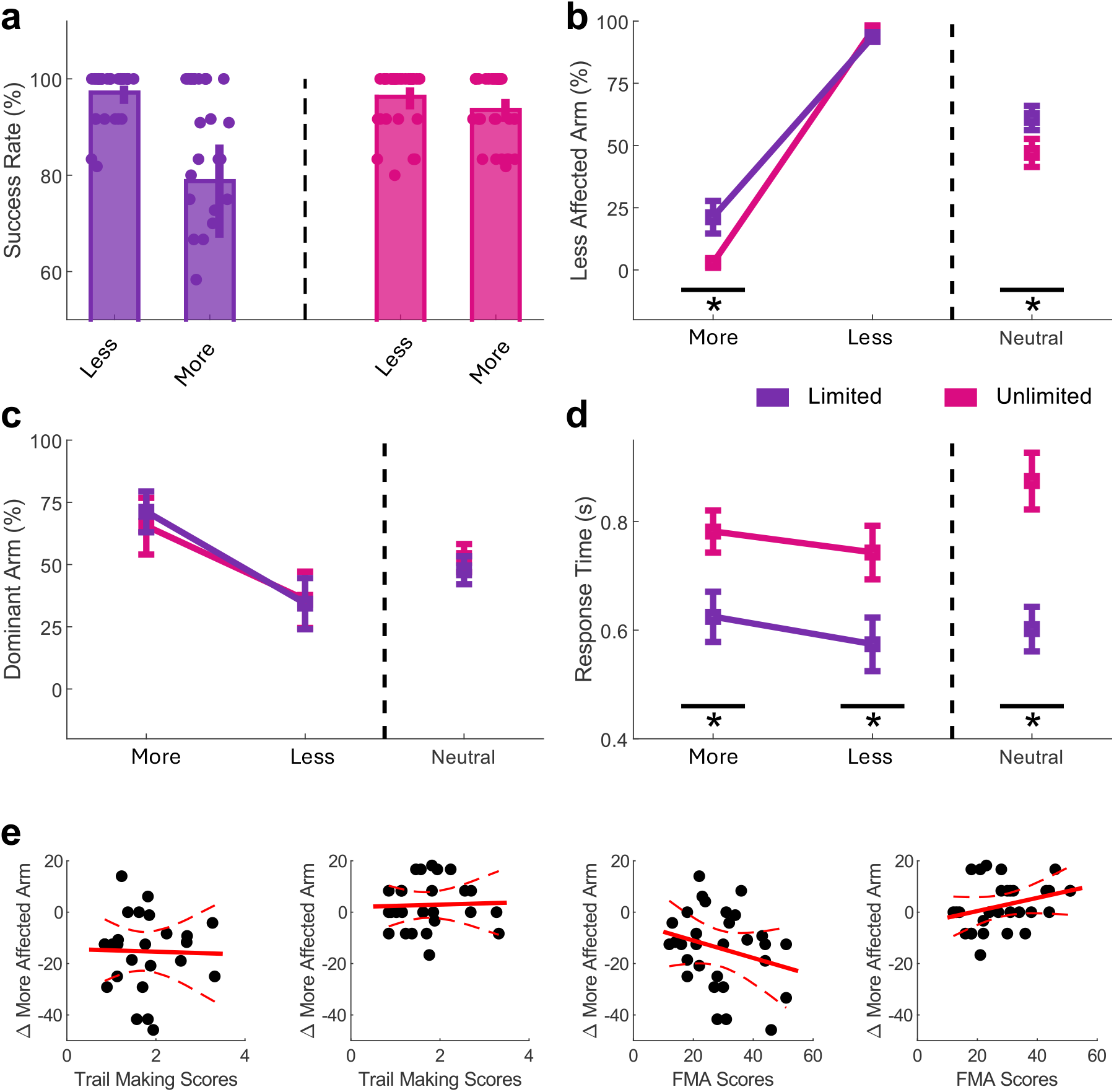
Non-use may reflect a habit response. **a)** *Success Rate* (%) decreases only in *the Paretic/Limited* condition suggesting the presence of a habit. **b)** Change in *Non-paretic Arm Use* (%) in each Choice Condition from the *Unlimited* and *Limited* Response Condition. In *Paretic* participants lose out on 10 points, because using the non-paretic arm was the less rewarding option. An increase across Response Condition indicates an increase in habitual behaviour. In contrast, in *Non-Paretic* participants win 10 points, because here using the non-paretic arm was the more rewarding option. Therefore, here habit and most rewarding option align. In *Neutral* using either arm yields the same reward. **c)** Change in *Dominant Arm Use* (%) (%) from the *Unlimited* and *Limited* response time condition for all Choice Conditions. **d)** Change in *Response Times (s)* from the *Unlimited* and *Limited* response time condition for all Choice Conditions. **e)** Left: Relationship between changes in the use of the paretic arm (%) and Trail Making scores (standardised z-scores) in the *Neutral* (right) and *Paretic* (left) conditions. Bottom row: Relationship between changes in the use of the paretic arm (%) and FMA scores (only including Elbow and Shoulder sub-scores) in the *Neutral* (right) and *Paretic* (left) conditions.

Specifically, on average participants used the non-paretic arm in only ∼2% of trials in the *Paretic/Unlimited* condition. This, however, increased to ∼22% of trials in the *Paretic/Limited* condition despite this yielding fewer points. In contrast, no changes could be observed in the *Non-Paretic* condition across the *Limited* and *Unlimited* ‘Response Condition’.

#### 2. Use of formerly dominant arm does not increase in the *Limited* condition

We did not find evidence that *Dominant Arm Use* changed across Response Conditions. The results from the repeated-measures ANOVA did not reveal a significant main effect for ‘Response Condition’ (F = 0.01, p = 0.9351, η^2^ < 0.01, Figure 2c) but for ‘Choice Condition’ (F = 477.21, p < 0.0001, η^2^ = 0.96). Importantly, we also did not find a significant interaction for ‘Response x Choice Condition’ (F = 0.01, p = 0.9137, η^2^ < 0.01). These results highlight that the habit response expressed during the *Limited* response condition is not related to former arm dominance.

#### 3. *Response Times* significantly decreased in the *Limited* response time condition

The results from the the repeated-measures ANOVA revealed a significant main effect for ‘Response Condition (F = 67.96, p < 0.0001, η^2^ < 0.32, Figure 2f) and for ‘Choice Condition’ (F = 827.83, p < 0.0001, η^2^ = 0.96). Interestingly, we also found a significant interaction for ‘Response x Choice Condition’ (F = 55.19, p < 0.0001, η^2^ < 0.28). Post-hoc pairwise comparisons revealed that *Response Time* was significantly lower in the *Paretic* condition (Z = 3.75, p < 0.0001, d = 0.94, Figure 1d), *Neutral* condition (Z = 4.70, p < 0.0001, d = 1.52), and also in the *Non-Paretic* condition (Z = -1.84, p = 0.0682, d = -0.51) during the *Limited* response time condition.

#### 4. Decrease in *Arm Choice* is not related to motor and/or cognitive impairment

To understand if increases in *Paretic Arm Use* are related to motor and/or cognitive impairment, we correlated changes in use of the paretic arm (*ΔArm Use*) in both the *Paretic* and *Neutral* condition with participants FMA scores (only the FMA Elbow and Shoulder sub-scores were included) and TMT scores. We did not find *ΔArm Use* to be related to FMA and TMT scores in the *Paretic* condition (FMA: ρ = 0.31, p = 0.09; TMT: ρ = 0.04, p = 0.86), nor in the *Neutral* condition (FMA: ρ = -0.25, p = 0.16 TMT: ρ = 0.03, p = 0.90 Figure 2e). These results highlight that *ΔArm Use* are independent from motor and/or cognitive impairment.

## Discussion

Our results revealed a striking pattern of increased use of the non-paretic arm, even when it was the less rewarding choice (*Paretic* trials). Importantly, no change was observed in the *Non-Paretic* trials which highlights that participants continued to use the non-paretic arm when it was the more rewarding option. These results suggest that the use of the non-paretic arm reflects a habit response.

Our findings provide new insight into the phenomenon of “non-use” in chronic stroke patients, specifically that habitual behaviour may underlie the continued prioritization of the non-paretic arm. Non-use refers to the persistent avoidance or limited use of the paretic (weaker) limb, despite the potential for recovery of function. After a stroke, many patients experience hemiparesis, a condition where one side of the body is weakened. Initially, reliance on the non-paretic arm is often necessary to achieve immediate functional goals such as dressing or feeding. However, this reliance often continues long after some degree of recovery has been achieved, a phenomenon referred to as “learned non-use” (Taub et al., 2006). Our results suggest that this non-use may be partly driven by habitual responses.

Habits are not always present in every action but exert a strong influence on behaviour when established. A habitual response is automatic and fast but tends to be inflexible, favouring actions that were previously rewarded. In contrast, goal-directed behaviour is more deliberate and flexible but slower to enact (Robbins and Costa, 2017). In our study, participants continued to favour the non-paretic arm, even when using the paretic arm would have been more rewarding. This suggests that their behaviours was driven more by habit than by goal-directed decision-making.

The absence of a difference in errors during the non-paretic trials supports this conclusion: participants were able to continue making the correct choice when using their non-paretic arm, indicating they could still perform goal-directed actions when the task aligned with their habitual tendencies. The increased errors during paretic trials, however, suggest that habitual non-use of the paretic arm interferes with performance when switching to goal-directed action is required.

Our findings align with previous work that has questioned the neurophysiological basis of non-use, suggesting instead that it may have a cognitive origin (Kwakkel et al., 2015). While some deficits in motor function may be directly tied to neurological damage, the persistence of non-use is often better explained by cognitive and behavioural factors—such as habit formation—rather than ongoing neurophysiological impairments. The evidence that non-use might be underpinned by habitual behaviour is particularly intriguing, given the substantial body of literature on habit formation, breaking, and modification in other domains, such as addiction and dieting (Wood and Rünger, 2016).

Habit-breaking strategies, such as context manipulation, deliberate practice, and reward restructuring, have been shown to successfully alter habitual behaviour in various contexts. These strategies could be readily adapted to stroke rehabilitation to address habitual non-use. For example, constraint-induced movement therapy (CIMT) already forces the use of the paretic arm, essentially breaking the habit of using the non-paretic limb. However, our findings suggest that other cognitive-behavioural strategies, such as modifying environmental cues or increasing awareness of habit-driven behaviours, could complement CIMT and enhance its effectiveness.

In conclusion, recognizing that habitual behaviour may contribute to non-use opens up new avenues for rehabilitation strategies. Drawing from extensive research on habit change in other areas could significantly advance stroke rehabilitation approaches, offering patients more tailored and effective interventions to restore functional use of the paretic arm.

## Acknowledgements

This work was supported by the Jon Moulton Charity Trust and the Stroke Association, UK.

